# Ulinastatin Ameliorates Cardiac Ischemia/Reperfusion Injury via Inhibiting the tissue kallikrein-kinin system

**DOI:** 10.1101/2023.02.21.529463

**Authors:** Xiao Ran, Hang Ruan, Shu-sheng Li, Rongxue Wu, Ailin Luo, Qin Zhang

## Abstract

**Background:** Endothelial cells (ECs) are considered more sensitive to cardiac ischemia/reperfusion (I/R) injury compared to cardiomyocytes. However, current research is mainly focused on molecular mechanisms and preventive strategies targeting cardiomyocyte I/R injury, whereas insufficient attention is placed on protecting endothelial function.

**Methods and Results:** In this study, we established an interlink among ulinastatin (UIT; a serine protease inhibitor), the kallikrein-kinin system (KKS), and EC injury in response to cardiac reperfusion for the first time, using in vitro and in vivo experiments, and bioinformatic analysis. Our data indicated that UTI affected I/R by inhibiting the activation of KKS and simultaneously down-regulating both bradykinin receptor 1 (Bdkrb1) and bradykinin receptor 2 (Bdkrb2) related signaling such as extracellularsignal–regulated kinase (ERK)/inducible nitric oxide synthase (iNOS) and vascular endothelial growth factor (VEGF)/endothelial nitric oxide synthase (eNOS), thereby reducing infarct size, attenuating inflammation and edema, and improving cardiac function and mortality.

Interestingly, UIT significantly suppressed KLK1 activity but did not down-regulate the KKS in normal conditions, suggesting inhibition of KLK1 might be the crucial mechanism for UIT-induced cardioprotection in reperfusion injury. Moreover, knockdown of Bdkrb1 in reperfusion-induced cardiac endothelial cells (MCECs) injury significantly prevented ERK translocation into the nucleus, reducing apoptosis, junction disruption, and expression levels of cytokines, whereas Bdkrb2 deletion could not protect MCECs against I/R injury.

**Conclusions:** Our findings imply that inhibition of KLK1/Bdkrb1 is a critical target for UIT in the treatment of reperfusion-induced cardiac endothelial inflammation, apoptosis, and leakage and might be a potential therapeutic strategy for cardiac reperfusion injury.

## Introduction

Ischemic heart disease remains the leading cause of morbidity and mortality worldwide^1^. Cardiac ischemia/reperfusion (I/R) injury is considered an important therapeutic target for ischemic heart disease^2^. Primary percutaneous coronary intervention and fibrinolytic therapy, which restore blood flow to the ischemic myocardium and limit infarct size, are the most common reperfusion therapies^3, 4^; yet, current research is mainly focused on molecular mechanisms and preventive strategies targeting cardiomyocyte I/R injury, whereas insufficient attention is placed on protecting cardiac cell function.

Studies have shown that compared with cardiac myocytes, endothelial cells (ECs) are less sensitive to ischemic injury but more sensitive to reperfusion injury in the heart^5^. In addition, continuous evidence supports the primary role of ECs in acute cardiac I/R injury-induced cardiomyocyte death^6^. Endothelial activation and injury increase vascular permeability, promote inflammatory cell invasion, and mediate myocardial damage after reperfusion^7^.

The kallikrein-kinin system (KKS), present on the vascular endothelium, has an essential role in maintaining the homeostasis of the cardiovascular system^8^. As a key enzyme of the KKS, tissue kallikrein (KLK1) is a serine protease that can cleave low-molecule kininogen into bradykinin and kallidin^9^. Under normal conditions, bradykinin activates endothelial nitric oxide synthase (eNOS) primarily via bradykinin receptor 2 (Bdkrb2) and produces temporary nitric oxide (NO), mediating cardiovascular protection. In response to I/R, bradykinin receptor 1 (Bdkrb1)-dependent inducible nitric oxide synthase (iNOS) is activated and releases a large amount of NO, mediating inflammation and increasing EC permeability^10, 11^. Considering the two receptors of KKS mediating different effects, our current understanding of the role of KKS in response to cardiac I/R injury is still rather limited. Thus, further exploring the mechanisms of KKS in cardiac I/R injury and identifying potential therapeutic targets are of utmost importance.

Ulinastatin (UTI), a serine protease inhibitor derived from human urine, was found to have anti-inflammatory and cytoprotective actions. It has been widely used in China, Japan, and Korea for treating patients with inflammatory disorders^12, 13^. UTI can suppress various serine proteases and protect EC against neutrophil-mediated injury^14^. Moreover, recent studies have found that UTI protects against I/R^15, 16^ and may be favorable for those patients undergoing cardiac surgery with cardiopulmonary bypass^17^. However, the current understanding of the role of UIT on cardiac I/R is limited.

Since KLK1 is a serine protease and an important element of KKS, we hypothesized that UIT exerts anti-inflammatory effects and improves endothelial barrier function by inhibiting the KKS activation in response to I/R. In order to identify KKS involved in UIT-induced cardioprotection after I/R, we screened the mRNAs for their differential expression levels in the heart of mouse cardiac I/R model and discovered that the potential protective effect of UIT on I/R might act by inhibiting KKS and simultaneously down-regulating both of Bdkrb1 and Bdkrb2 related signaling, such as ERK/iNOS and VEGF/eNOS. We then examined the role of UIT in the treatment of cardiac I/R injury and explored the underlying molecular mechanism of KKS in the UIT treatment of cardiac reperfusion injury following ischemia. Specifically, we isolated primary mouse cardiac endothelial cells (MCECs) to establish an interlink among UIT, KKS, and EC injury in response to cardiac reperfusion and found that UIT protects the heart from I/R injury by suppressing the activation of KKS and then reducing inflammation and junction disruption. Our findings imply that UIT treatment and inhibition of KLK1/Bdkrb1 signaling are potential therapeutic strategies for cardiac reperfusion injury.

## Materials and methods

### Animals

Male adult mice (8-week-old) were purchased from the Hunan SJA Laboratory Animal Co., Ltd (Changsha, Hunan, China). Mice were housed in a facility with a 12-h light/12-h dark cycle at 22-24 °C and received water and food *ad libitum*. Animal care and experimental procedures were carried out in accordance with the guidelines provided by the Institutional Animal Care and Use Committee of Tongji Hospital, Tongji Medical College, Huazhong University of Science and Technology (Wuhan, China).

### Mouse model of cardiac I/R injury

Left anterior descending (LAD) coronary artery ligation was used to induce I/R in mice, as previously described^18^. Briefly, the animals were anesthetized with isoflurane (3% isoflurane for induction, 2% isoflurane for maintenance). They were then orally intubated with a 22G IV catheter and artificially ventilated with a rodent respirator (DW-3000A/B, tidal volume 0.3ml, rate 105 strokes/min). Next, the left of the chest was cut open to expose the heart, and the LAD coronary artery was ligated via a slipknot using 6–0 silk for 45 min, after which the slipknot was gently loosened to induce reperfusion injury. Mice in the sham group underwent the same procedures without tying the slipknot. Then, sixty mice were randomly divided into three groups: Sham, I/R + vehicle, and I/R + UIT. Before reperfusion, the mice injected with a total volume of 200ul saline were set as I/R + vehicle group, and an equal amount of UIT (15 U/g, Techpool Bio-Pharma Co., Ltd. Guangzhou, China) was given to mice by the tail vein in I/R +UIT group.

### Echocardiography

The cardiac function was analyzed 24 hours after I/R using transthoracic echocardiography, as previously reported^19^. Mice were anesthetized with inhalation of isoflurane (1 to 1.5%) in O2, and their chest hair was removed using depilatory paste. M-mode echocardiography was performed with a VEVO 770 system (VisualSonics, Toronto, Canada). The short-axis and long-axis views were obtained to measure the cardiac parameters. The percentage of ejection fraction (EF) and fractional shortening (FS) was detected.

### Evans Blue extravasation assay

Evans blue was used to evaluate the permeability of the heart as previously described ^20^. Mice were restrained so that their tail was accessible. After the vein was located, the needle was inserted by directing the needle into the vein with its bevel pointing upward at an angle of approximately 20 degrees. The needle was slowly inserted, visualizing the needle as it entered the vein. Once the vein’s wall had been penetrated, the needle was cranially directed approximately 2 mm. Blood was then aspirated into the needle’s hub before making an injection. Then, mice were injected intravenously with 200 μL of 0.5% Evans Blue dye. (Cat No. E2129; Sigma Aldrich). Consequently, the pressure was applied over the injection site by gently holding a piece of gauze over the injection site for approximately 30 seconds to prevent hematoma formation. Thirty minutes after the injection, mice were euthanized, and their hearts were harvested for further experiments.

### Triphenyltetrazolium chloride (TTC) staining

The area of infarction was determined by 2, 3, 5-triphenyltetrazolium chloride (TTC) staining in I/R hearts^18^. After 24 hours of reperfusion, the thread around the LAD was re-ligated, and 0.5 mL of 1% Evans blue solution was reversely injected into the aortic root while the aorta was clipped with a hemostat. Apart from the anterior descending branch area, other non-ischemic areas of the heart were stained. The heart was then extracted and sectioned. The sections were subsequently incubated with 1% TTC (CatNo.1779; Sigma Aldrich) for 15 min at 37 °C, and photographs were obtained with a digital camera. The blue part represented normal non-ischemic myocardial tissue (ANAR), the red part represented the area at risk (AAR), and the white part represented myocardial tissue with ischemic infarct (INF). The Image J software was employed to measure the blue, red, and white areas of the section and calculate the percentage of ischemic risk area and myocardial infarct area.

### Measurement of the myocardial water content

Heart samples were weighed and dried in a thermostat oven at 80°C for 48 h. The myocardial water content was calculated according to the following formula: myocardial water content (%) = [(wet weight - dry weight) / wet weight] × 100%.

### Histological and IHC staining

After 72 hours of reperfusion, the mice were euthanized, and the hearts were harvested. The heart tissues were washed with PBS, fixed for 1 hour at room temperature in 4% paraformaldehyde solution, and perfused with 30% sucrose at 4°C overnight before being embedded in the optimal cutting temperature (OCT). Heart sections were then stained with hematoxylin and eosin (H&E, Cat No.G1120; Solarbio) and Masson trichrome (Masson, Cat No.G1340; Solarbio) staining following the manufacturer’s protocol. For IHC staining, paraffin-embedded hearts were cut transversely into 5-μm sections. Next, sections were incubated with the primary antibodies, anti-Bdkrb1 (CatNo.G1340; Invitrogen), in 3 % BSA in PBS overnight at 4 °C followed by respective secondary antibodies (KIT-5020; MaxVisionTM HRP-Polymer anti- Mouse/Rabbit IHC Kit). Images were captured with Leica Application Suite X software (Leica).

### TUNEL assay

Heart sections (described above) were used for TUNEL assays using the in situ cell death detection kit (Cat No. 11684817910, Roche), according to the manufacturer’s instructions. Nuclei were stained with DAPI (4′,6-diamidino-2-phenylindole). Images were acquired using a fluorescence microscope (DM1000, Leica Microsystems, Germany).

### Serum collection and measurement

Blood was collected from tail veins 24h after the operation. Samples were immediately centrifuged, after which the supernatant was collected and stored at -80°C. Serum bradykinin (Cat No. MU30950; Bioswamp, China), IL-1β (Cat No. MU30369; Bioswamp, China), IL-6 (Cat No. MU30044; Bioswamp, China) and TNF-α (Cat No. MU30030; Bioswamp, China) levels were measured using ELISA kits, respectively.

### Determination of KLK1 activity and NO Production in heart tissue

Hearts collected from mice 24 hours after I/R were homogenized, and the supernatant was used for KLK1 activity detection. Next, an assay utilizing the immunological characteristics of TK and the catalysis of benzoyl arginine ethyl ester hydrochloride (BAEE) was used to determine KLK1 activity^21^. Finally, NO production in heart tissue was analyzed by Total Nitric Oxide Assay Kit (Cat No. S0023; Beyotime, China).

### RNA Sequencing

RNA sequencing data were collected from 9 samples of 3 groups (Sham, I/R + vehicle, and I/R +UIT). To identify differentially expressed genes (DEGs) related to I/R and UTI treatment, R package limma^22^ was used to determine DEGs between every two groups, which implements an empirical Bayesian approach to estimate gene-expression changes using moderated t-tests. DEGs were determined by significance criteria (adjusted P value < 0.05) as implemented in the R package limma. The intersection of these DEGs was visualized with Vennplot. Heatmaps were conducted to reveal the differential expressions of these genes. To identify the potential function and involved pathways related to UTI treatment in I/R, we conducted Gene Ontology (GO) and Kyoto Encyclopedia of Genes and Genomes (KEGG) analyses based on the differential expression profiles between every two groups, using the clusterProfiler^23^ R package. Bubble plot and concept-gene plot were used to display the pathways that these DEGs were enriched in.

### Exploring Key Genes Interrelated with UTI Treatment Process

To find out the key genes involved in the UTI treatment process, we selected the DEGs that were up-regulated in the IR group and Sham group and down-regulated in the IR-UTI group and IR group. A total of ninety-four genes and their expression profiles were visualized by heatmap. STRING^24^ was used to analyze the protein interaction network (PPI) among these genes, and the top 10 core genes in the network were filtered out. The expression profiles of the 12 key genes in three groups were displayed with a violin plot.

### The immunological characteristics of the immune cell microenvironment (TME) in three groups

The Cibersort^24^ algorithm was used to quantify the relative abundance of 21 immune cells in the TME based on specific immune cell gene sets. The relationship between the relative abundance of 21 immune cells and the expression matrix of the 12 key genes was calculated and visualized by corrplot, and the 3 most related immune cells’ relative abundance was visualized with violin plot.

### Isolation of cardiac endothelial cells from mice

Isolation and purification of endothelial cells from mouse heat were described previously^25, 26^, which used mechanical and enzymatic dissociation followed by activated magnetic cell sorting for purification. Briefly, mouse hearts were removed from 6-8 weeks mice and then digested with 2 mg/ml collagenase (CatNo.LS004196; Worthington). After digestion, the supernatant was harvested and centrifuged to collect the pellet. Then, the pellet was re-suspended in a solution containing PBS without calcium and magnesium (GIBCO), BSA (Bovine Serum Albumins) (0.1% v/v, Sigma), and platelet endothelial cell adhesion molecule-1 (PECAM1, CD31) coated beads (CatNo.BDB553370; Fisher). After separating cells from the magnetic separator, cells with beads were plated in a gelatin-coated T25 flask. About 5-9 days when cells approached and became confluence, the second sort with an identical protocol to the above was performed using intercellular adhesion molecule 2 (ICAM2, CD102) coated beads (CatNo.BDB553325; Fisher).

### siRNA transfection and quantitative RT-PCR (qRT-PCR)

siRNA duplexes were designed, synthesized, and purified by RiboBio Co., Ltd (Guangzhou, China). MCECs were seeded in triplicate in tissue culture dishes (24-well or 96-well) 24 h prior to siRNA transfection. Control, Bdkrb1, or Bdkrb2 siRNAs were transfected at a concentration of 50 nM using Lipofectamine RNAiMAX (Invitrogen) according to the manufacturer’s instructions. The efficiency of siRNA for knockdown was assessed by qRT-PCR. siRNA sequences and primers for qRT-PCR are listed in **Table S1**. Total RNA was isolated from cells subjected to real-time PCR (Applied Biosystems). Quantitative PCR analysis was conducted on the CFX-Connect 96 system (Bio-Rad) with SYBR Green PCR Master Mix (Cat No. KM4101; KAPA Biosystems) using validated primers specific to each target of each gene.

### Determination of apoptosis

Quantification of apoptotic MCECs was determined by flow cytometry-based Annexin V-FITC/PI staining (CatNo.556547, BD Bioscience) according to the manufacturer’s protocol, followed by flow cytometry analysis.

### Antibodies

The following antibodies were used for WB and immunofluorescence IF: rabbit Anti-VE-Cadherin (CatNo.361900, Life Technologies; 1:1000 for WB and 1:200 for IF), mouse Anti-ZO-1 (CatNo.339100, Invitrogen; 1:1000 for WB and 1:200 for IF), rabbit Anti- Bdkrb1 (CatNo.577292, Invitrogen; 1:1000 for WB and 1:200 for IF), rabbit Anti- Bdkrb2 (CatNo.550488, Invitrogen; 1:1000 for WB and 1:200 for IF), rabbit Anti- iNOS (CatNo.30111, Bioswamp; 1:1000 for WB and 1:200 for IF), rabbit Anti- iNOS (CatNo.30111, Bioswamp; 1:1000 for WB and 1:200 for IF), mouse Anti-eNOS (CatNo.ab76198, Abcam; 1:1000 for WB and 1:200 for IF), rabbit Anti-VEGF (CatNo.ab52917, Abcam; 1:1000 for WB and 1:200 for IF), rabbit Anti- IL-1β (CatNo. 37329, Bioswamp; 1:1000 for WB and 1:200 for IF), rabbit Anti- IL-6 (CatNo. 37326, Bioswamp; 1:1000 for WB and 1:200 for IF) and rabbit Anti- TNF-α (CatNo. 37416, Bioswamp; 1:1000 for WB and 1:200 for IF).

### Transendothelial electrical resistance (TEER) measurement

TEER was measured using an automated impedance sensing system (ECIS; Applied Biophysics, Troy, NY) as previously described^27^. MCECs were seeded at a density of 60,000 cells/cm^2^ in each well of the manufacturer’s eight-well electrode slide (8W10E). ECIS was conducted using the multiple frequency/time (MFT) options to record the impedance measurements over a broad spectrum of frequencies.

### Statistical analysis

All analyses were analyzed with the R (version 4.1.0) and R Bioconductor packages. The chi-square (χ2) test was performed to evaluate differences between the two groups. Normally distributed continuous data are presented as mean ± SE and compared using Student’s t-test, while non-parametric data were presented as median (interquartile range) and compared using Mann–Whitney U test as appropriate. The Spearman correlation coefficient was used to evaluate correlations. The Kaplan–Meier survival curve was analyzed with the chi‐squared test. A two-sided p-value <0.05 was considered statistically significant for all tests.

## Results

### UIT protects against cardiac I/R Injury in mice

We hypothesized that UIT might reduce cardiac damage in response to reperfusion injury since UIT can improve cardiac blood vessel barrier function, alleviate cardiac edema formation, and protect against heart failure^15, 16^. First, we determined the cardioprotective effect of UIT *in vivo* in a mouse acute cardiac I/R model with 45 minutes of ischemia followed by 24 hours of reperfusion (**Figure 1A**) and found that the infarct size relative to the area at risk (INF/AAR) in mice treatment of UIT (before the onset of reperfusion) decreased from 71.0±3.1% to 23.5±3.0%, concomitantly with similar AARs (**Figure 1B-D**).

**Figure 1.**
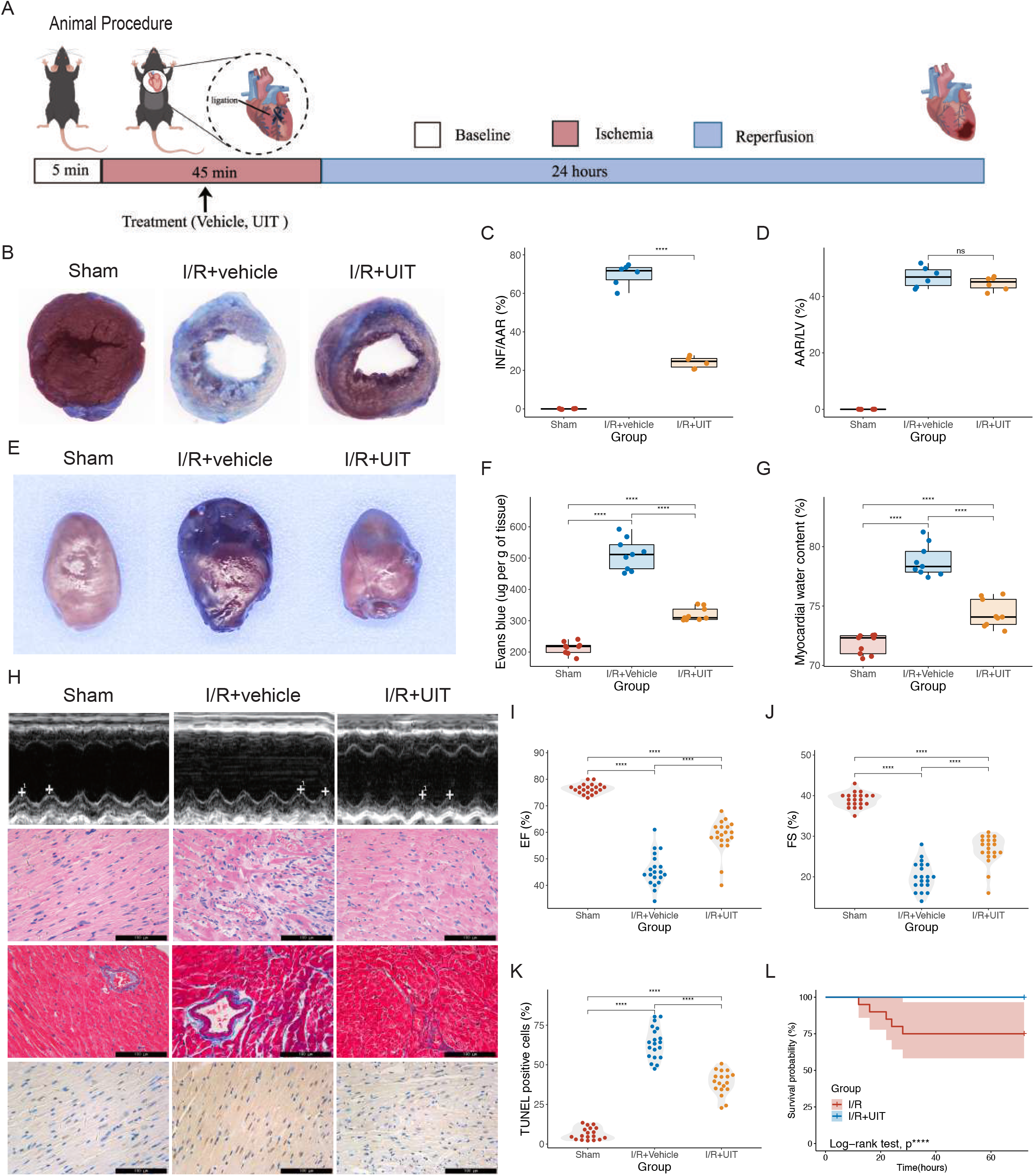
Ulinastatin (UIT) protects against cardiac ischemia-reperfusion (I/R) injury in mice. **(A)** Experimental protocol showing duration and time course of ischemia (I) and reperfusion (R). **(B)** Representative photographs of heart sections and quantified data demonstrate that UIT decreased infarct size in mice subjected to ischemia/reperfusion (I/R, 45 min/24 h). **(C, D)** A myocardial area at risk (AAR) as a percentage of the total left ventricle (LV) and infarct size (INF) as a percentage of AAR evaluated at 24 h post-reperfusion (n=6 mice in each group). **(E)** Representative images of Evans Blue leakages after I/R injury 24 hours. **(F)** Quantification of Evans Blue extravasation in cardiac tissue from different groups following I/R injury (n=9 mice in each group). **(G)** Myocardial water content measured in different groups after I/R 24 hours (n=9 mice in each group). **(H)** Representative images of echocardiographic images, H&E (×400 magnification), Masson (×400 magnification), and TUNEL staining (×400 magnification) 24 hours after reperfusion. **(I-J)** Fractional shortening (FS) and ejection fraction (EF) were analyzed by echocardiography (n=20 mice in each group). **(K)** Histogram analysis of the cell apoptosis (%) by Image J software (n = 18 in each group). **(L)** Kaplan–Meier survival curves. All the mice were randomly divided into three groups: Sham, I/R + vehicle, and I/R + UIT. Before reperfusion, I/R + vehicle group was injected with a total volume of 200ul saline by tail vein, while the I/R +UIT group received an equal amount of UIT. Data shown as means ± SEM, and statistical significance was determined using two-tailed t-test or two-tailed log-rank test (****p<0.0001).

Next, we evaluated endothelial barrier integrity and cardiac edema by EB leakage and water content. As expected, increased leakage of Evans blue dye and water content were detected in the vehicle-treated groups receiving I/R injury, while treatment with UIT remarkably decreased EB extravasation in the heart and attenuated cardiac edema (**Figure 1E-G**). Echocardiography showed that UIT improves I/R-induced cardiac dysfunction, as indicated by the increased ejection fraction and fractional shortening (**Figure 1H-J**). In addition, histological staining, including H&E and Masson staining, showed treatment with UIT at the beginning of reperfusion largely attenuated I/R-induced myocardial interstitial edema, rupture of myocardial fibers, infiltration of leukocytes and collagen deposition (**Figure 1H**). Also, a significant decrease in TUNEL-positive cells was detected in mice treated with UIT compared with vehicle-treated mice in the acute cardiac I/R model, indicating that UIT decreases myocardial apoptosis after I/R injury (**Figure 1H, K**). Furthermore, a comparison of Kaplan-Meier survival curves showed a lower mortality rate in UIT-treated I/R mice compared to the vehicle-treated I/R mice (**Figure 1 L**). Taken together, these data suggest that UIT promotes cardioprotection from acute cardiac I/R injury.

### Inhibition of KKS is a key mechanism in UIT-induced cardioprotection after I/R

The DEGs between the groups of Sham and I/R are shown in **Figure 2**. Compared with Sham, 2926 differential expressed genes were found in the I/R group (**Figures 2C**) and 5265 in the I/R-UTI group. There were 1740 differential expressed genes in the I/R-UTI group compared to the I/R group (**Figures 2C-E**). The intersection of the three kinds of DEGs was visualized by vennplot (**Figure 2F**). Heatmaps were conducted to visualize the intersection of all three kinds of DEGs (**Figure 2G**), while GO and KEGG analyses were conducted based on differentially expressed genes from different groups. The DEGs’ pathways are shown in a bubble plot in **Figure S1**, and the genes involved with these pathways are shown in **Figure S2**.

**Figure 2.**
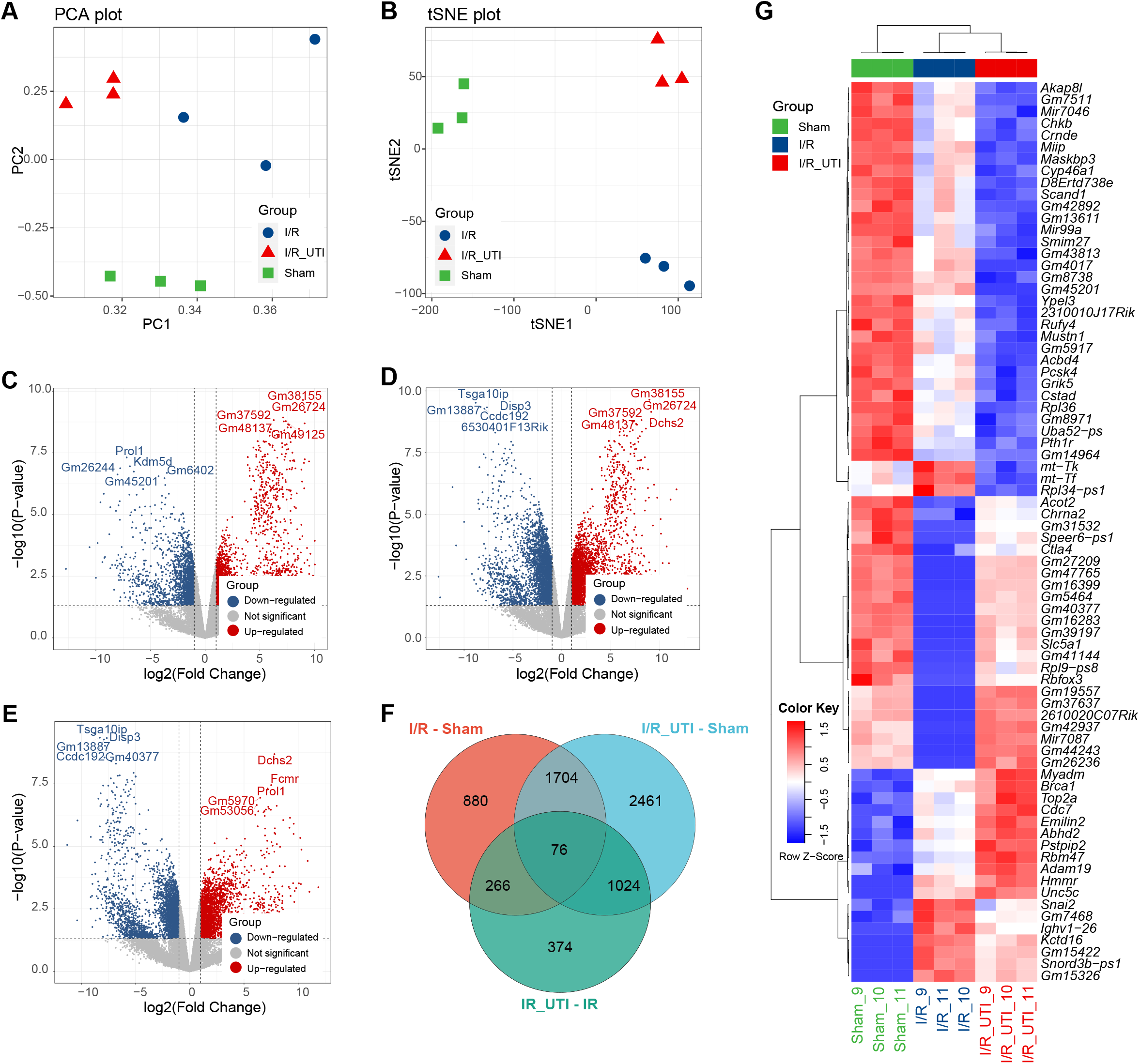
The expression characteristics in 9 samples of 3 groups. **(A)** PCA dimension reduction algorithm was conducted to evaluate the differences between 3 groups. **(B)** tSNE dimension reduction algorithm was conducted to evaluate the differences between 3 groups. **(C)** The differential expressing genes of the total RNA expression profile between I/R samples and Sham samples was visualized by Vioplot. **(D)** The differential expressing genes of the total RNA expression profile between I/R-UTI samples and Sham samples was visualized by Vioplot. **(E)** The differential expressing genes of the total RNA expression profile between I/R samples and I/R-UTI samples was visualized by Vioplot. **(F)** Venn plot showed the intersection of the 3 kinds of DEGs. **(G)** Heatmaps presented the expression of 76 intersection differential expression genes of the 3 kinds of DEGs.

Results of bioinformatics analysis found that 94 genes were up-regulated in the I/R group and down-regulated in the I/R-UTI group (**Figure 3A**). Most of them significantly interacted with each other, which shows an inner connection among these genes **(Figure 3B, C**). KLK1, the key enzyme of KKS, was identified as an important UIT-treatment-related gene in I/R. As a receptor of KKS, Bdkrb1 was also identified as a key gene in UIT-treated I/R mice and correlated with KLK1 in the present study. Further analysis considered that 12 genes related to KKS were interrelated with the UTI treatment process, and their expression in the three groups is shown in **Figure 4**. These suggested that the potential protective effect of UIT on I/R was regulated by inhibiting KKS and simultaneously down-regulating both Bdkrb1- and Bdkrb2-related signalings, such as MAPK/iNOS and VEGF/eNOS. We concluded that UIT alleviated reperfusion-induced inflammation and cardiac blood vessel barrier function by reducing the expression of proinflammatory cytokines (IL-1β, IL-6, and TNF-α) and increasing junction levels (ZO-1 and VE-cadherin) via inhibiting the Bdkrb1/iNOS signal pathway and down-regulating the expression of VEGF via Bdkrb2/eNOS signaling.

**Figure 3.**
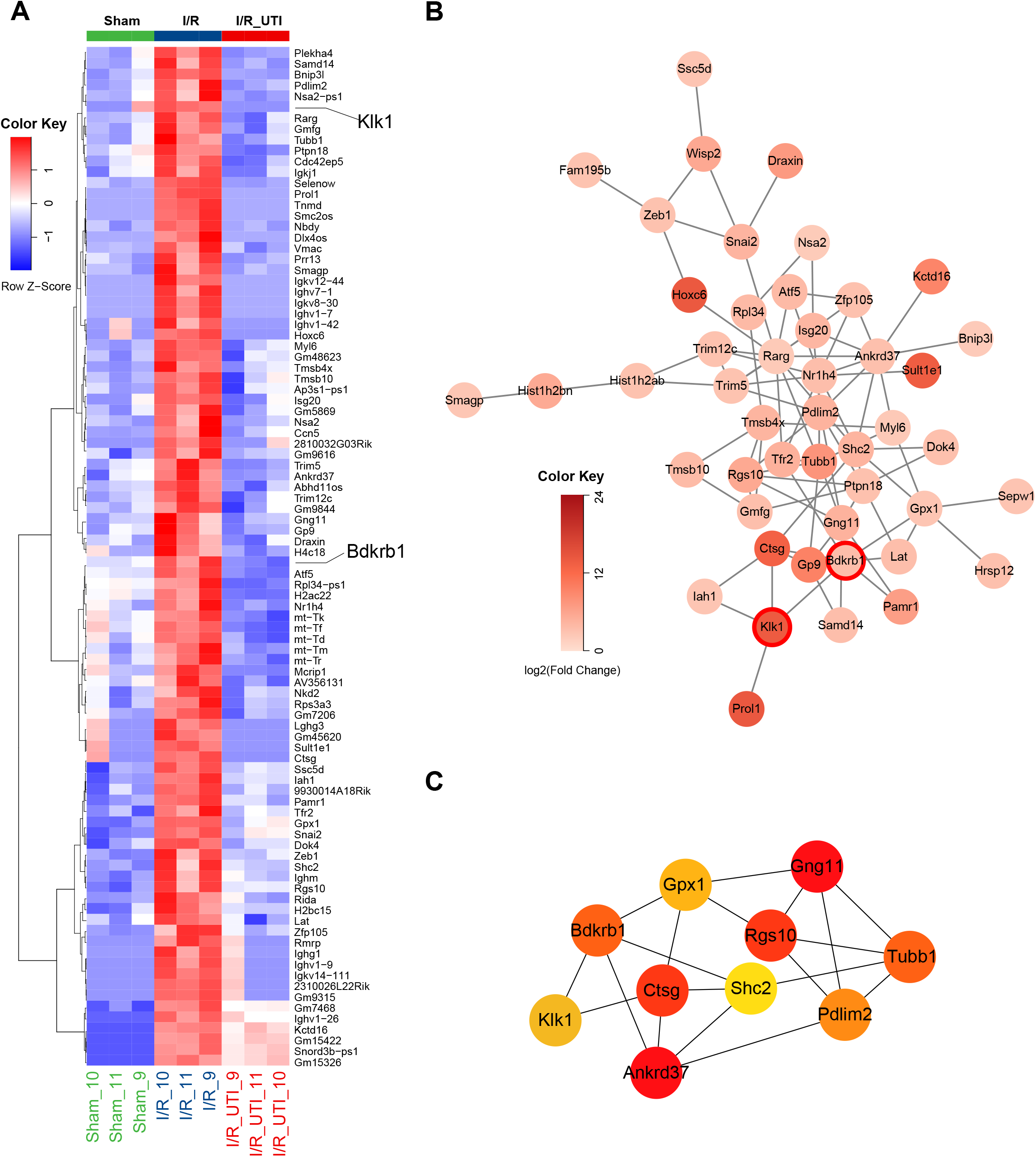
The expression features of the up-regulated genes in I/R groups. **(A)** Heatmaps presented the expression of 94 up-regulated genes in I/R groups. **(B)** The protein-protein interaction network of the 94 genes. **(C)** Top 10 core genes in the protein-protein interaction network.

**Figure 4.**
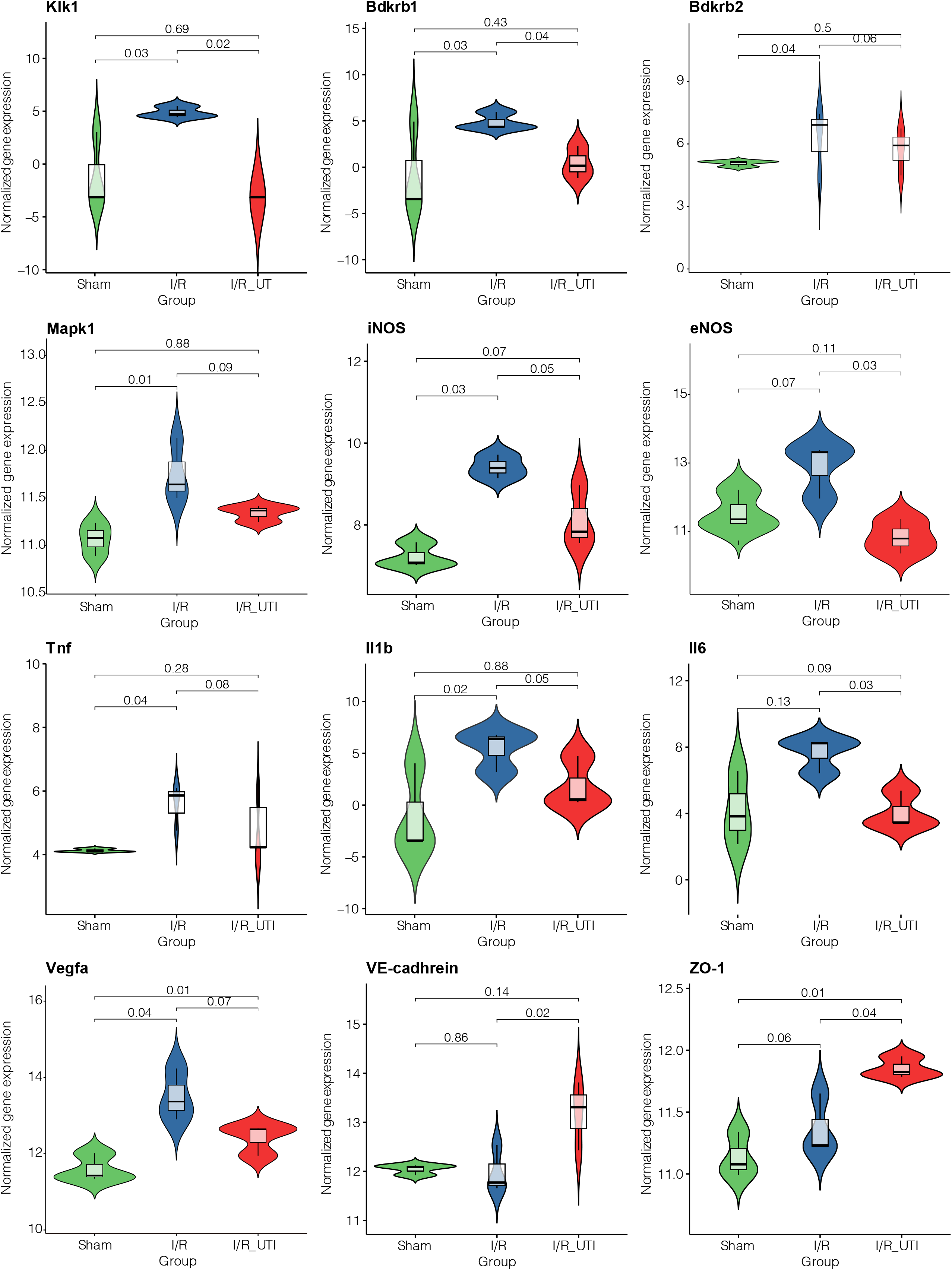
Differences in expression features of the 12 UTI treatment-related genes in different groups.

### Depicting Immunological Characteristics and the Relationship between Immune Cells and Key Genes

Next, we explored the immune microenvironment in the 9 samples and quantified the concentration of immune cells with the Cibersort algorithm. As shown in **Figure 5A**, M1 macrophage showed significantly high levels in I/R samples compared to Sham and I/R-UTI samples, which was also demonstrated by the violin plot (**Figure 5C**). Also, M1 macrophage showed a relationship with most UTI treatment-related genes (**Figure 5B**). In addition, neutrophils, plasma cells, T CD8 Naive cells, and Th17 cells were also related to some of the key genes.

**Figure 5.**
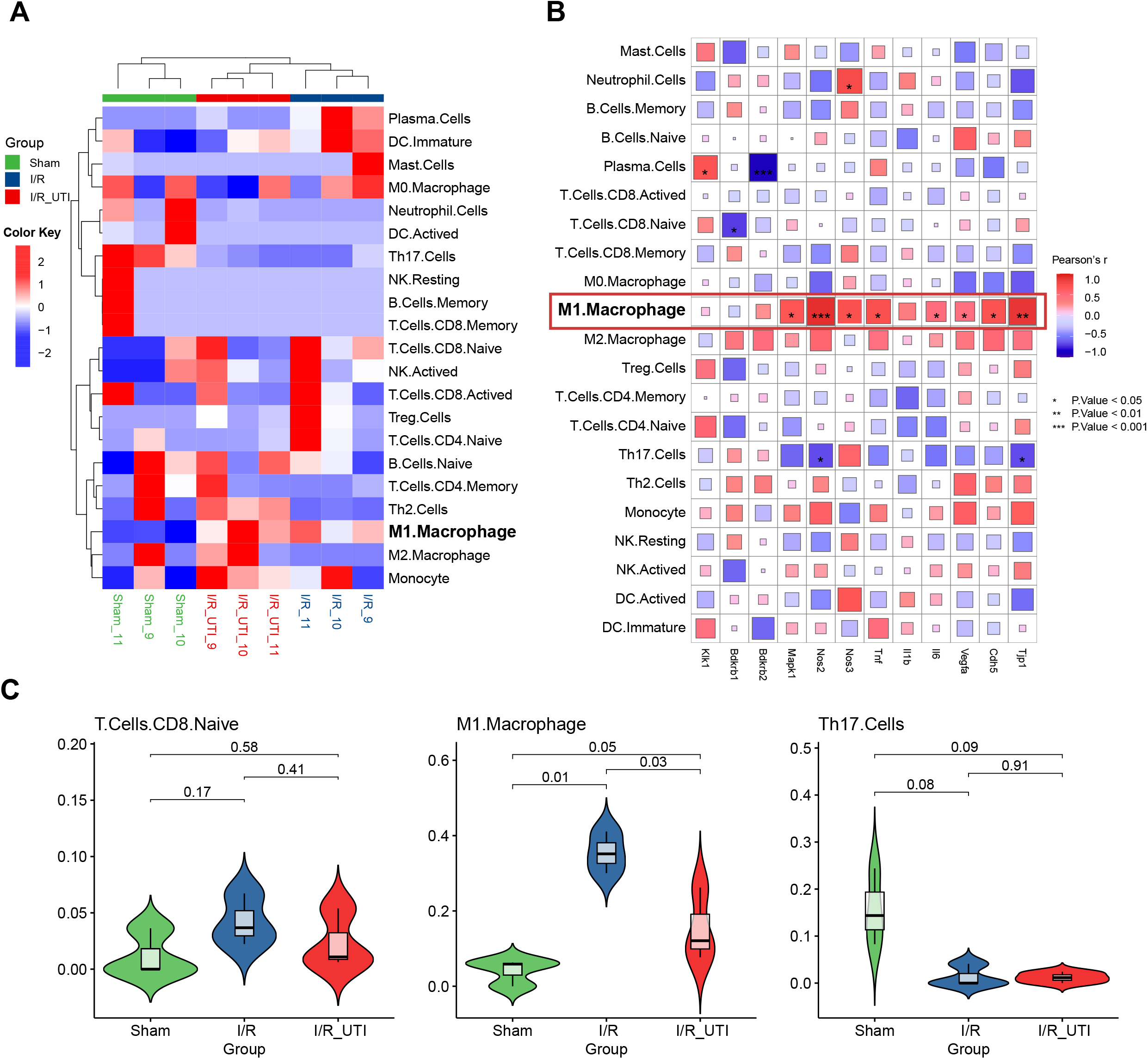
The immunological characteristics of the samples and their relationship with UTI treatment-related genes. **(A)** Differences in immune cells among three groups. **(B)** The correlation of the immune cells and the UTI treatment-related genes was visualized by corrplot. **(C)** The differences between the three most significant immune cells in the three groups.

### UIT alleviates cardiac reperfusion injury by inhibiting KKS-mediated inflammation and junction disruption

To validate our bioinformatic predictions, the role of UIT on KKS-mediated cytokines (IL-1β, IL-6, and TNF-α), junctions (ZO-1 and VE-cadherin), and VEGF in response to I/R was investigated *in vivo*. KLK1, as a key enzyme for the KKS, regulates the end-effector substance bradykinin through the corresponding receptors (Bdkrb1 and Bdkrb2) and then mediates iNOS and eNOS activation and NO production. KLK1 activity (**Figure 6A**), expression levels of Bdkrb1/iNOS and Bdkrb2/eNOS (**Figure 6G-H** and **Figure 7A-E**), and NO production (**Figure 6C**) in heart tissue and plasma bradykinin levels (**Figure 6B**) increased significantly after 24 h of I/R, suggesting the KKS was up-regulated in response to I/R. This was accompanied by a sharp rise of plasma levels of IL-1β (345±42 versus 121±9 pg/ml, *p*<0.001; n=5; **Figure 6D**), IL-6 (706±60 versus 91 ± 10 pg/ml, *p*<0.001; n=5; **Figure 6E**) and TNF-α (398 ± 31 versus 100±8 pg/ml, *p*<0.001; n=5; **Figure 6F**) and protein levels of cytokines (**Figure 7F-H**), junctions (**Figure7I-J**), and VEGF (**Figure 7K**). Moreover, results of biochemistry, ELISA, immunohistochemical staining, and Western blotting suggested that all members of the KKS in the heart tissue present much lower levels in UIT-treated I/R mice compared to the vehicle-treated I/R group (**Figure 6A-C** and **Figure 7A-E**), as well as KKS-mediated cytokines (**Figure 6D-F** and **Figure 7F-H**), junctions (ZO-1 and VE-cadherin), and VEGF in heart tissue and plasma. However, it is interesting that UIT could not down-regulate expression levels of the KKS and related proteins except for KLK1 in the Sham group. These results suggested that inhibition of KLK1 might be the crucial mechanism for UIT-induced cardioprotection in reperfusion injury.

**Figure 6.**
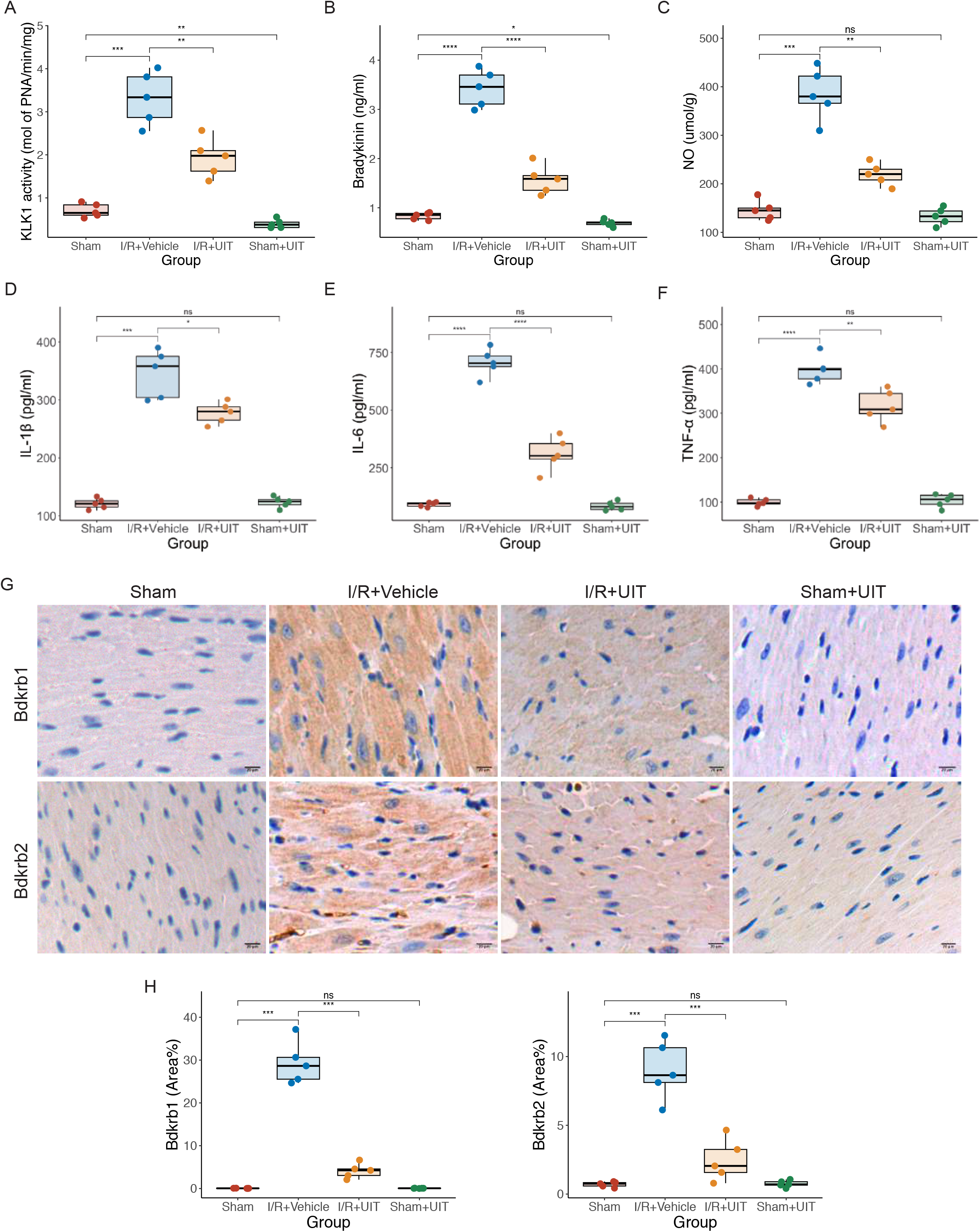
Ulinastatin (UIT) suppresses cardiac reperfusion-induced elevated levels of the KKS and cytokines in mice. **(A)** The activity of KLK1 in the heart tissue of mice. **(B-C)** Protein levels of bradykinin and NO in the heart tissue of mice. **(D-F)** Serum IL-1β, IL-6, and TNF-α levels in mice. **(G)** Immunohistochemical staining for Bdkrb1 and Bdkrb2 (magnification, ×400). **(H)** Immunohistochemical analysis of Bdkrb1 and Bdkrb2. N=5 mice in each group. Data shown as means ± SEM, and statistical significance was determined using a two-tailed t-test (*p<0.05, ** p<0.01; *** p<0.001****p<0.0001).

**Figure 7.**
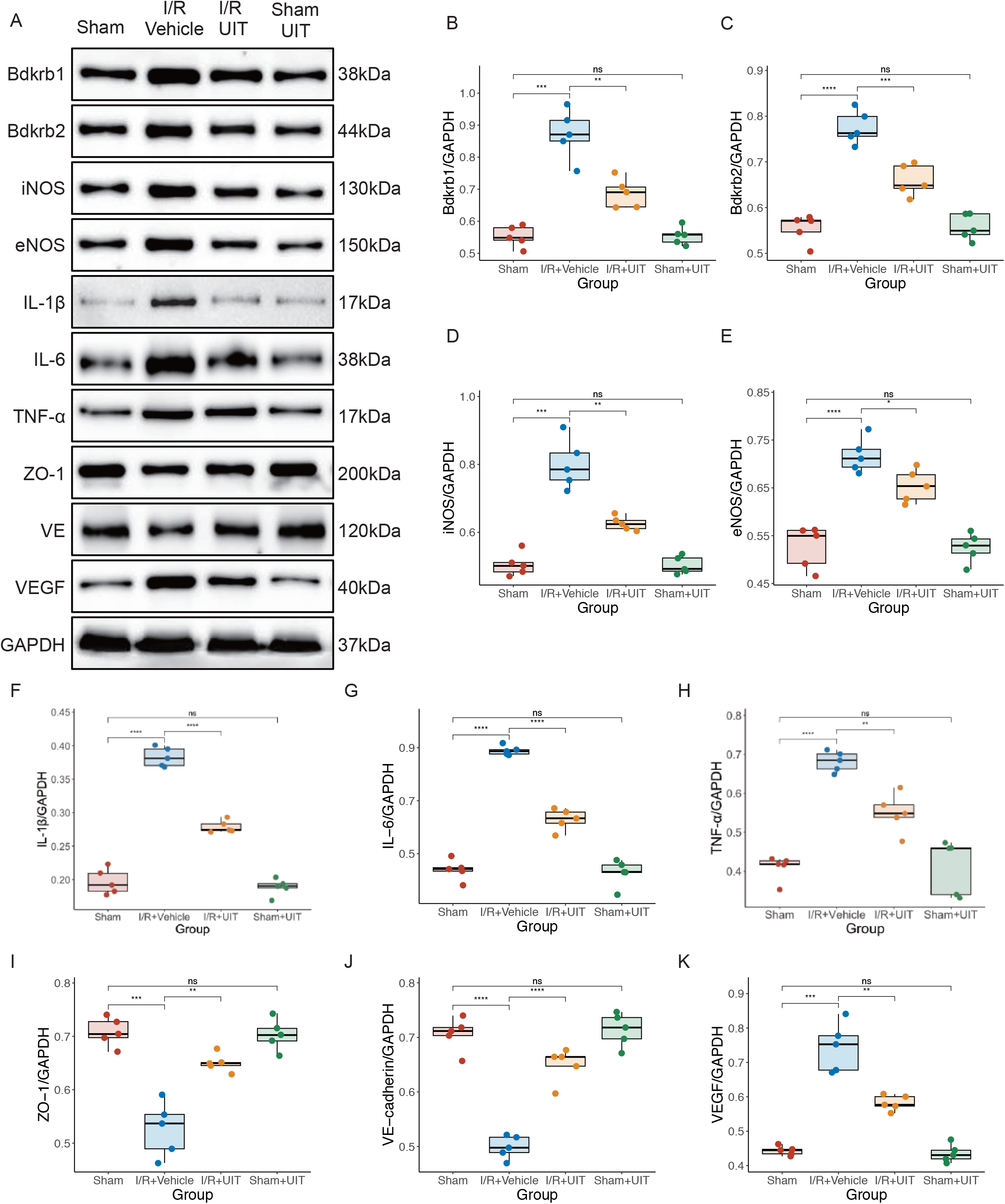
Ulinastatin (UIT) inhibits KKS-mediated inflammation and junction disruption in the heart tissue of mice after cardiac reperfusion. **(A)** Representative western blot of Bdkrb1, Bdkrb2, iNOS, eNOS, IL-1β, IL-6, TNF-α, ZO-1, VE-cadherin and VEGF. **(B-K)** Western blot analysis of Bdkrb1, Bdkrb2, iNOS, eNOS, IL-1β, IL-6, TNF-α, ZO-1, VE-cadhrein and VEGF (n=5). Data shown as means ± SEM, and statistical significance was determined using a two-tailed t-test (*p<0.05, ** p<0.01; *** p<0.001****p<0.0001).

### UIT reduces the activation of KKS, inflammation, and junction disruption caused by reperfusion injury in cardiac endothelial cells

Our bioinformatic analysis and *in vivo* experiments suggested that UIT alleviated cardiac reperfusion injury by inhibiting KKS-mediated inflammation and junction disruption. The KKS exerts physiological effects by binding kinin peptides to the Bdkrb1 and Bdkrb2, which are localized in endothelial cells. Oxygen-glucose deprivation-reoxygenation (OGD/R) is a widely accepted model for studying ischemic reperfusion *in vitro*. Thus, we performed a series of *in* vitro experiments to validate our findings from the bioinformatic analysis and *in vivo* study. In the supernatant of primary MCECs cultured, KLK1 activity markedly increased after OGD/R (**Figure 8A**). Meanwhile, the expression of BK receptors (Bdkrb1 and Bdkrb2) (**Figure 8B-F**) and their related molecules (iNOS and eNOS), cytokines (IL-1β, IL-6, and TNF-α), and VEGF in MCECs were up-regulated significantly in response to OGD/R, while protein levels of ZO-1 and VE-cadherin were down-regulated (**Figure 8D-N**). When MCECs were treated with UIT at the onset of reperfusion, the results were consistent with our bioinformatic analysis and *in vivo* experiments; the levels of KKS (KLK1, Bdkrb1, Bdkrb2, iNOS, and eNOS), cytokines (IL-1β, IL-6, and TNF-α) and VEGF decreased, and protein levels of junctions (ZO-1 and VE-cadherin) were significantly increased in MCECs (**Figure 8B-N**). These results suggested that UIT might alleviate reperfusion-induced inflammation and junction disruption via inhibition of KKS in cardiac endothelial cells.

**Figure 8.**
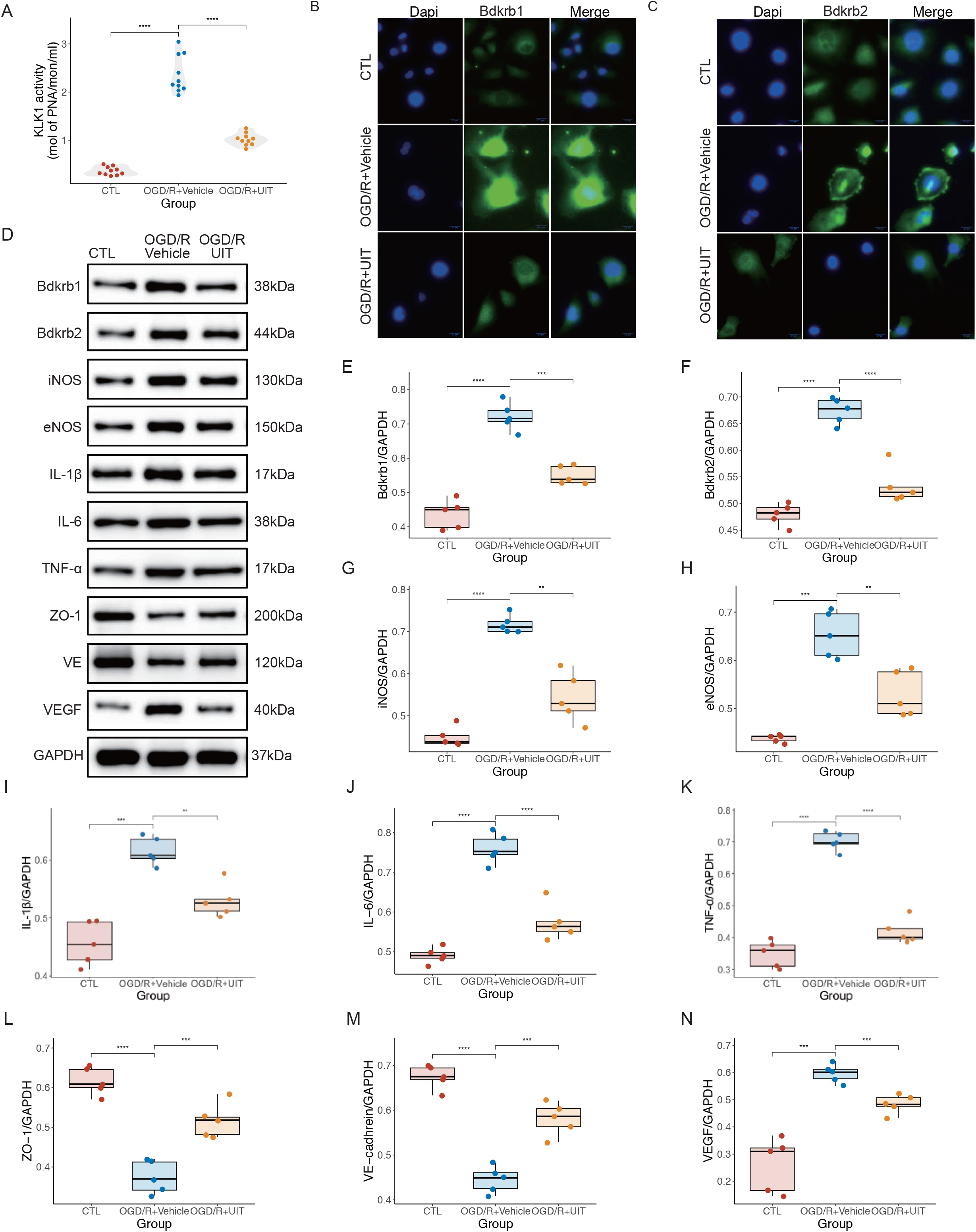
Ulinastatin (UIT) reduces the activation of KKS, inflammation, and junction disruption caused by reperfusion injury in cardiac endothelial cells (MCECs). **(A)** KLK1 activity in extracts from MCECs (n=10). **(B, C)** Immunofluorescence staining of Bdkrb1 and Bdkrb2 in MCECs. **(D)** Representative western blot of Bdkrb1, Bdkrb2, iNOS, eNOS, IL-1β, IL-6, TNF, ZO-1, VE-cadherin, and VEGF. **(E-N)** Western blot analysis of Bdkrb1, Bdkrb2, iNOS, eNOS, IL-1β, IL-6, TNF-α, ZO-1, VE-cadhrein and VEGF (n=5). Data shown as means ± SEM, and statistical significance was determined using a two-tailed t-test (** p<0.01; *** p<0.001****p<0.0001).

### Bdkrb1 is a critical target for UIT in the treatment of reperfusion-induced cardiac endothelial inflammation, apoptosis, and leakage

As most of the physiological activities induced by kinins are mediated by BK receptors (Bdkrb1 and Bdkrb2) belonging to the G-protein-coupled receptor family, we knockdown these receptors by corresponding siRNA and explored their role on reperfusion-induced cardiac endothelial leakage and inflammation. siRNA knockdown was confirmed by real-time PCR. Knockdown of *Bdkrb1* and *Bdkrb2* significantly reduced the expression of Bdkrb1/iNOS and Bdkrb2/eNOS. Also, reperfusion-induced elevated levels of Bdkrb1/iNOS and Bdkrb2/eNOS decreased by corresponding siRNA and UIT (**Figure 9A-D**). These were further confirmed by Western blot analysis (**Figure 9E-I**). Interestingly, elevated cytokines levels (IL-1β, IL-6, and TNF-α) induced by OGD/R could be sharply decreased by *Bdkrb1* siRNA and UIT instead of *Bdkrb2* siRNA (**Figure 9J-L**), suggesting Bdkrb1 deletion and treatment with UIT could reverse OGD/R-induced inflammation.

**Figure 9.**
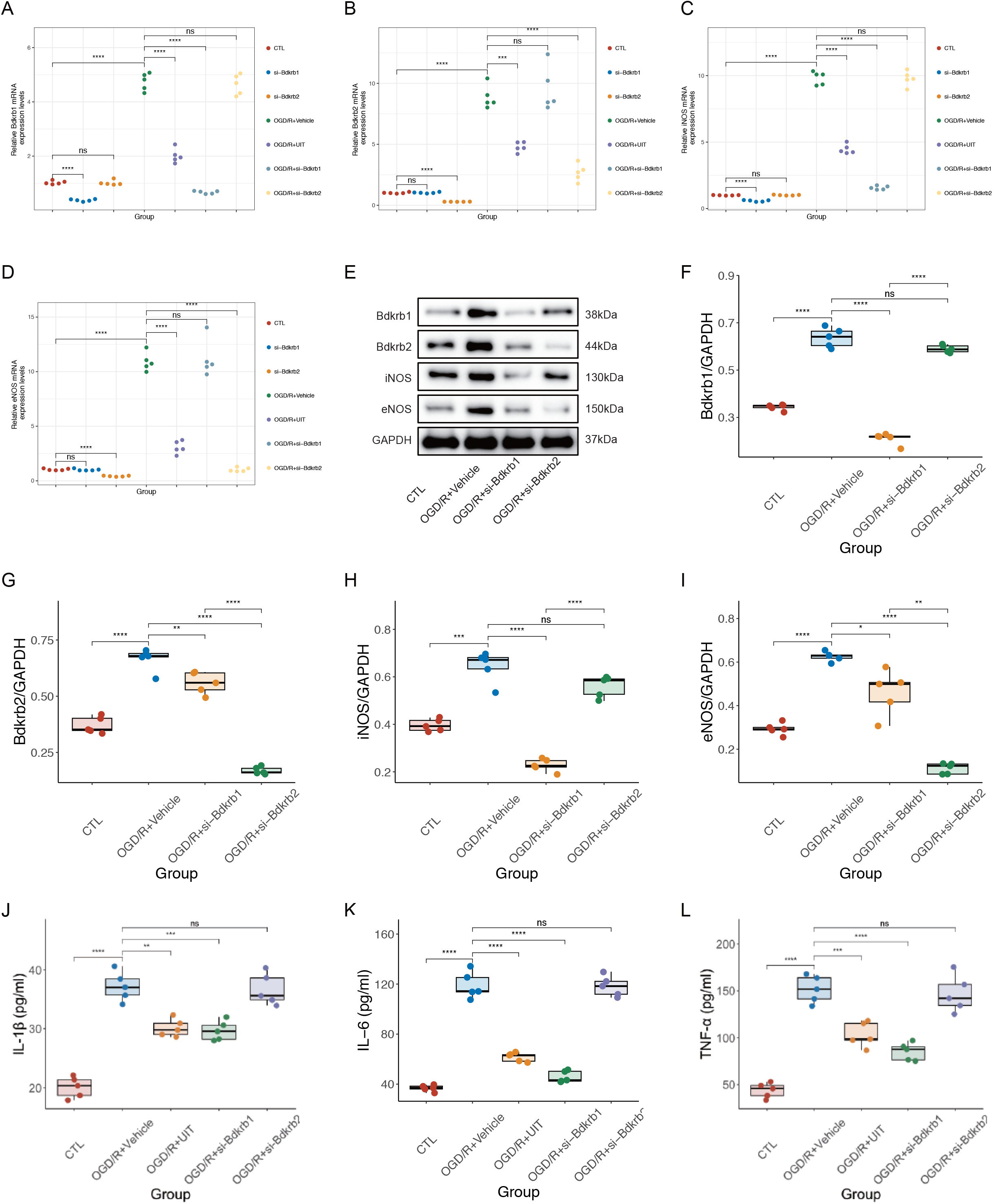
Bdkrb1 deletion reverse OGD/R-induced inflammation in cardiac endothelial cells (MCECs). **(A-D)** mRNA levels of Bdkrb1, Bdkrb2, iNOS and eNOS in MCECs (n=5). **(E)** Representative western blot of Bdkrb1, Bdkrb2, iNOS, and eNOS. (F-I) Western blot analysis of Bdkrb1, Bdkrb2, iNOS, and eNOS (n=5). **(J-L)** Protein levels of IL-1β, IL-6, and TNF-α in cell culture medium (n=5). Data shown as means ± SEM, and statistical significance was determined using a two-tailed t-test (** p<0.01; *** p<0.001****p<0.0001).

To further verify the anti-apoptosis role of UIT we observed *in vivo*, we measured apoptosis in cardiac endothelial cells in the OGD/R model and found that UIT treatment and deletion of Bdkrb1 markedly decreased OGD/R-induced apoptosis. As shown in **Figure 10A**, the percentage of apoptotic cells rose to nearly 28% after OGD/R insult when MCECs were treated with vehicle and *Bdkrb2* siRNA, while it was significantly decreased in the UIT and Bdkrb1 siRNA-treated OGD/R groups. Therefore, these results, combined with our *in vivo* study, suggest that Bdkrb1 deletion inhibits OGD/R-induced apoptosis.

**Figure 10.**
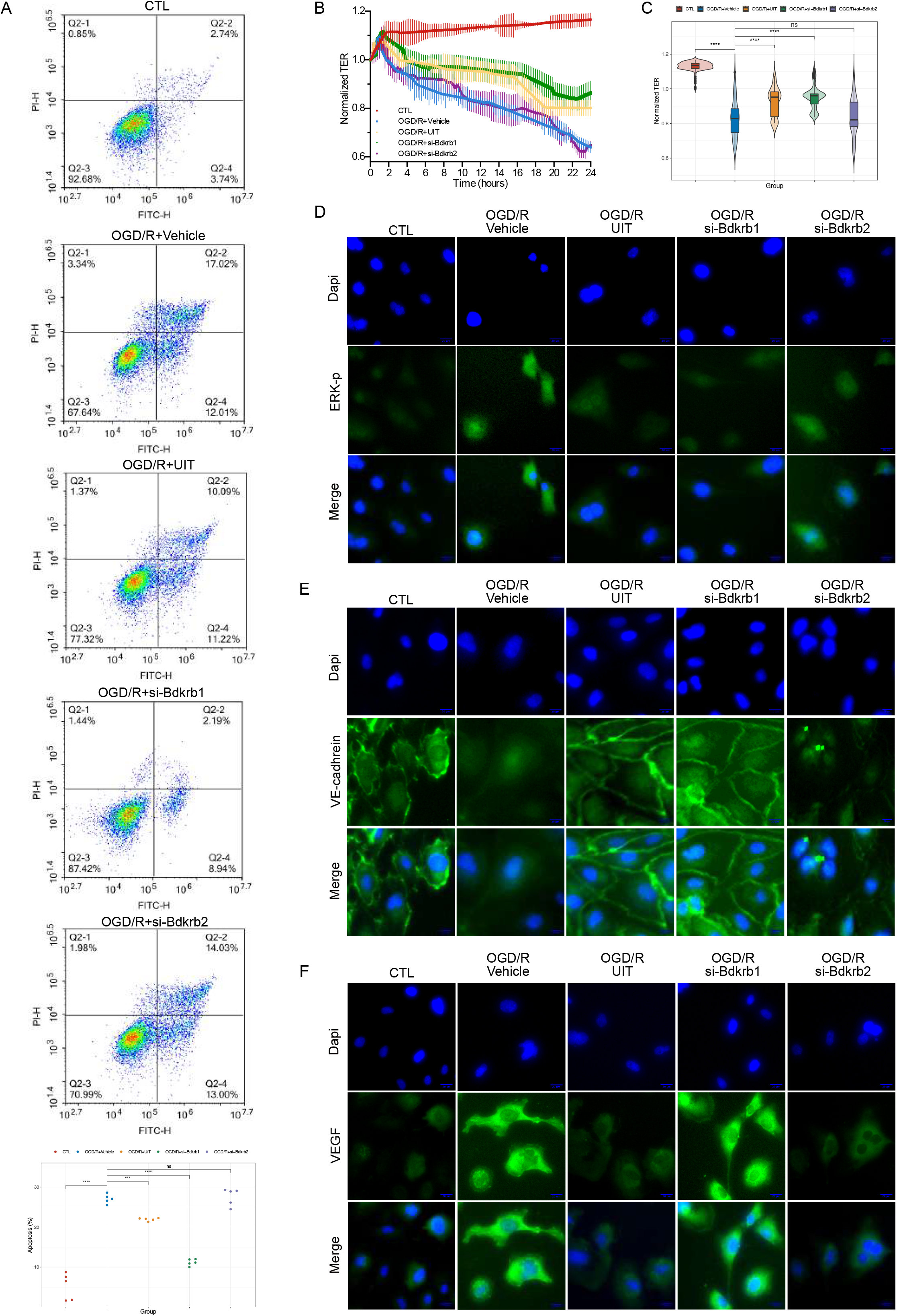
Inhibition of Bdkrb1 and treatment of ulinastatin (UIT) suppress reperfusion-induced apoptosis and hyperpermeability in cardiac endothelial cells (MCECs). **(A)** The percentage of apoptotic cells was determined by flow cytometric analysis (n=5). **(B)** Trans-endothelial electrical resistance (TER) across confluent MCECs was monitored over time with an electrical cell-substrate impedance-sensing system. **(C)** Mean relative the endothelial barrier resistance detected by ECIS. Data shown as means ± SEM, and statistical significance was determined using a two-tailed t-test (*** p<0.001; ****p<0.0001). **(D-F)** Immunofluorescence staining of ERK-p, VE-cadherin, and VEGF.

Next, we measured changes in endothelial resistance on *in vitro* reperfusion using the electrical cell-substrate impedance sensing system (ECIS) (Applied BioPhysics, NY). Because transendothelial electrical resistance (TEER) measurements are inversely correlated with membrane permeability, the significantly lower measurements were observed in OGD/R-treated MCECs than MCECs that had been treated with vehicle, indicating that OGD/R significantly increased MCEC permeability. Resistance measurements in UIT- or *Bdkrb1* siRNA-treated MCECs also declined when cells were cotreated with OGD/R but were significantly higher than in OGD/R-treated MCECs. However, there was no difference in resistance measurements between *Bdkrb2* siRNA- and vehicle-treated MCECs when cells were treated with OGD/R. These data suggest that inhibition of Bdkrb1 and treatment of UIT could considerably suppress reperfusion-induced cardiac endothelial hyperpermeability.

Consistent with our speculation, IF staining showed that ERK-p was significantly up-regulated in the nucleus while endothelial junctional marker (VE-cadherin) along the borders of the disrupted area was dramatically down-regulated when MCECs were treated with OGD/R. Notably, cotreatment with UIT or *Bdkrb1* siRNA remarkably counteracted OGD/R-induced elevated expression of ERK-p and disrupted VE-cadherin at the cell boundary in MCECs; nevertheless, cotreatment with *Bdkrb2* siRNA did not appreciably alter the expression of ERK-p and VE-cadherin in OGD/R treated MCECs. *Bdkrb2* siRNA, but not UIT and *Bdkrb1* siRNA, could reverse OGD/R-induced increasing expression of VEGF in MCECs. Moreover, results in the present study suggested that UIT reduced the activation of KKS, including *Bdkrb1 and Bdkrb2*. Overall, these data indicated that Bdkrb1 was a critical target for UIT in the treatment of reperfusion-induced cardiac endothelial inflammation, apoptosis, and leakage.

## Discussion

Herein, we sought to test the hypothesis that UIT treatment could improve cardiac blood vessel barrier function, alleviate cardiac edema formation and inflammation, and protect against cardiac dysfunction after I/R. After treating mice with UIT at the beginning of reperfusion, a marked reduction in infarct size, infiltration of leukocytes, collagen deposition, and cardiac edema was seen. To further reveal the underlying mechanisms in the present study, RNA-seq analysis combined with *in vivo* and *in vitro* studies was conducted, suggesting that the potential protective effect of UIT on I/R acted by inhibiting KKS and simultaneously down-regulating both Bdkrb1 and Bdkrb2-related signalings such as ERK/iNOS and VEGF/eNOS. However, we also found that UIT could not down-regulate KKS except for KLK1 in normal conditions. Moreover, we systematically demonstrated for the first time that Bdkrb1 deletion remarkably suppressed reperfusion-induced cardiac endothelial inflammation, apoptosis, and leakage, whereas inhibition of Bdkrb2 could not. Together, our results highlight that KLK1/Bdkrb1 is a crucial target for UIT in the treatment of cardiac I/R injury (Figure 11).

**Figure 11.**
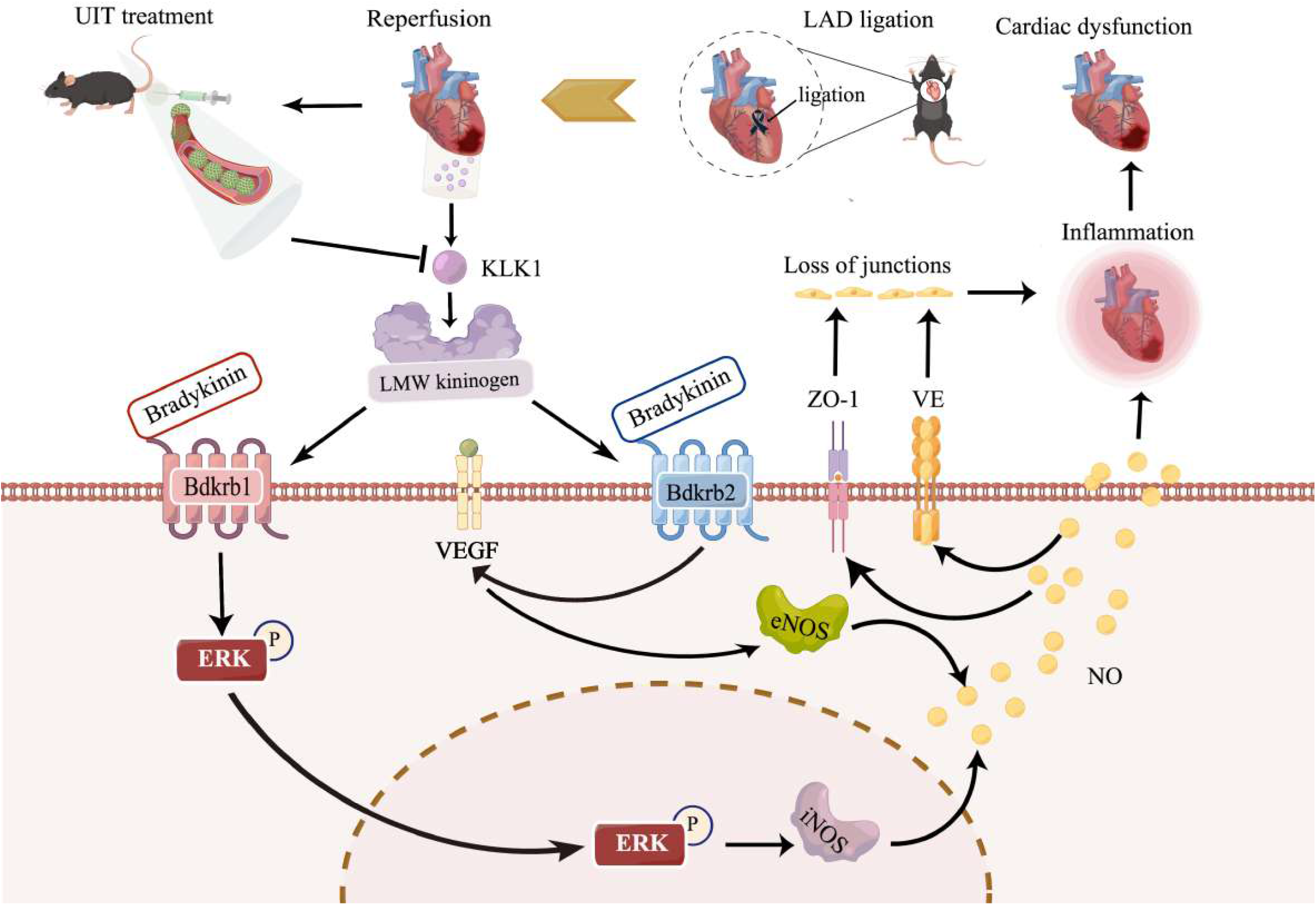
Mechanistic diagram summarizing the cardioprotective effect of ulinastatin (UIT) on ischemia-reperfusion (I/R).

Our study revealed a novel mechanism centered on KLK1 that connects UIT treatment and the KKS-related signaling regulation in cardiac inflammation and endothelial leakage after I/R injury, suggesting that inhibition of KLK1 could be the critical mechanism for UIT-induced cardioprotection in reperfusion injury. KLK1 is a serine protease and a key enzyme of the KKS^28^, while UIT is originally purified as a serine protease inhibitor^29^. Both of them can be extracted from healthy human urine^30, 31^. Thus, we considered that UIT’s role in treating I/R injury might be exerted by reducing KLK1 activity. As expected, KLK1 activity in the heart tissue was strongly suppressed in both the sham group and vehicle-treated I/R group. However, the global KKS-related signaling was dramatically increased in response to cardiac reperfusion, whereas it significantly decreased after UIT treatment (given before cardiac reperfusion). This is probably because cardiac reperfusion injury activates KLK1, activating the KKS cascade and then promoting inflammation, cardiac vascular permeability, and edema. Subsequently, the expression of Bdkrb1 and Bdkrb2, which mediate KKS-related signaling activation, was strongly up-regulated under inflammatory conditions^32-34^. Our previous study demonstrated that KLK1 was associated with inflammation and hyperpermeability of EC via high Bdkrb1 expression levels^31, 35^. Therefore, we supposed that treating mice with UIT before reperfusion could substantially inhibit KLK1 activity and suppress KKS-mediated inflammation and junction disruption.

This newly discovered connection between KKS-related signaling and cardiac I/R injury may provide an alternative perspective for numerous published studies. MAPK, VEGF, and KKS are highly expressed in response to I/R^11, 36-38^. Analysis of i*n vivo* RNA-Seq data combined with *in vivo* and *in vitro* studies suggest that UIT may reverse cardiac reperfusion-induced activation of MAPK, VEGF, and KKS-related signaling and then reduce the expression of proinflammatory cytokines and increase junction levels. We also discovered that KKS activates the KLK1/Bdkrb1/ERK pathway leading to increased levels of phosphorylated ERK in the nucleus and regulating the expression of iNOS and cytokines such as IL-1β, IL-6, and TNF-α in response to cardiac reperfusion, which, in turn, results in cardiac inflammation and disruption of the cardiac endothelial barrier. Also, cardiac reperfusion induces the activation of the KLK1/Bdkrb2/VEGF/eNOS signaling pathway, enhancing the cardiac vascular endothelial leakage. Thus, UIT can considerably suppress reperfusion-induced cardiac inflammation and endothelial hyperpermeability, inhibiting KKS-related signaling.

Our study indicated that elevated expression of iNOS is a major downstream event of Bdkrb1, mediating the activations of ERK after I/R and mediating cardiac inflammation, apoptosis, and leakage, which is consistent with the notion that “high-output” NO predominately produced by iNOS is a crucial indicator of inflammation and I/R^39, 40^. Indeed, our study revealed that Bdkrb1 deletion dramatically suppresses reperfusion-induced cardiac endothelial injury. Knockdown of Bdkrb1 in reperfusion-induced MCECs injury prevents ERK translocation into the nucleus and greatly diminishes apoptosis, junction disruption, and expression levels of cytokines. Although Bdkrb2 deletion could decrease the expression level of VEGF, our experiments suggest that Bdkrb2 knockdown could not protect the heart against I/R injury. Combined with *in vivo* results, these data indicate that Bdkrb1 mediating iNOS/NO signaling is the critical mechanism of cardiac I/R-induced injury. Therefore, we hypothesize that abolishing Bdkrb1 has great promise for cardiac reperfusion treatment. Hopefully, an endothelial-specific Bdkrb1 knockout mouse model will be able to further address the role of Bdkrb1 in cardiac I/R-induced injury in the future.

A direct connection between EC injury and myocardial dysfunction was reported in several studies^41, 42^. ECs lie near cardiomyocytes (CMs) in the heart and are considered uniquely situated to modulate CM function by synthesizing and releasing a variety of signaling molecules. They also might be a critical mediator of I/R injury in the heart^43^. Consistent with this notion, our data indicated that UIT suppresses the reperfusion-induced activation of KKS and then reduces inflammation and junction disruption in cardiac endothelial cells, consequently protecting CMs from apoptosis, inflammation, and edema, thereby improving myocardial function in our rodent model of I/R. Therefore, we considered that UIT improving cardiac function in cardiac I/R is mainly because of suppressing the KKS-induced disruption of endothelial barrier function and thus alleviating myocardial edema. It is also possible that UIT inhibits KKS-related inflammatory signaling in ECs in response to I/R, which may protect the myocardium against apoptosis and inflammation. However, the present study only focused on MCECs *in vitro*. Hence, further co-culture models of ECs/CMs are required to validate our results.

Our study suggested that the mechanism through which UIT treats cardiac I/R involves inhibiting KLK1 activity and then suppressing KKS-mediated inflammation and junction disruption. However, a limitation of the present study is that we could not disclose the therapeutic role of UIT in targeting KLK1, although our *in vivo* study demonstrated that UIT apparently decreased KLK1 activity of mice heart in normal conditions but could not down-regulate KKS and their related molecules. Therefore, *in vivo* studies such as endothelial-specific KLK1 knock-in mice will be used to demonstrate the cardioprotective effects of UIT in cardiac reperfusion is on-target inhibition of KLK1. Further in vitro studies, such as luciferase-reporter assays and chromatin immunoprecipitation (ChIP) analyses, will also be conducted to reveal the underlying molecular mechanisms.

To sum up, our data suggest that UIT regulates cardiac reperfusion by inhibiting KKS and simultaneously down-regulating Bdkrb1- and Bdkrb2-related signaling, such as ERK/iNOS and VEGF/eNOS, and reveals that inhibition of KLK1/Bdkrb1 is a crucial target for the therapy after cardiac I/R injury.

## Acknowledgements

This study was supported by funding from National Natural Science Foundation of China (82271358), the scientific Research Foundation for Returned Overseas Chinese Scholars of Tongji Hospital, Tongji Medical College, Huazhong University of Science and Technology, and Talents Project of Public Health in Hubei Province (2022SCZ048).

## Disclosures

The authors declare that they have no known competing financial interests or personal relationships that could have appeared to influence the work reported in this paper.

## Notes

### Competing Interest Statement

The authors have declared no competing interest.

